# Therapeutic efficacy of an oral nucleoside analog of remdesivir against SARS-CoV-2 pathogenesis in mice

**DOI:** 10.1101/2021.09.13.460111

**Authors:** Alexandra Schäfer, David R. Martinez, John J. Won, Fernando R. Moreira, Ariane J. Brown, Kendra L. Gully, Rao Kalla, Kwon Chun, Venice Du Pont, Darius Babusis, Jennifer Tang, Eisuke Murakami, Raju Subramanian, Kimberly T Barrett, Blake J. Bleier, Roy Bannister, Joy Y. Feng, John P. Bilello, Tomas Cihlar, Richard L. Mackman, Stephanie A. Montgomery, Ralph S. Baric, Timothy P. Sheahan

**Author notes:** These authors contributed equally to this manuscript (Alexandra Schäfer, David R. Martinez).

## Abstract

The COVID-19 pandemic remains uncontrolled despite the rapid rollout of safe and effective SARS-CoV-2 vaccines, underscoring the need to develop highly effective antivirals. In the setting of waning immunity from infection and vaccination, breakthrough infections are becoming increasingly common and treatment options remain limited. Additionally, the emergence of SARS-CoV-2 variants of concern with their potential to escape therapeutic monoclonal antibodies emphasizes the need to develop second-generation oral antivirals targeting highly conserved viral proteins that can be rapidly deployed to outpatients. Here, we demonstrate the in vitro antiviral activity and in vivo therapeutic efficacy of GS-621763, an orally bioavailable prodrug of GS-441524, the parental nucleoside of remdesivir, which targets the highly conserved RNA-dependent RNA polymerase. GS-621763 exhibited significant antiviral activity in lung cell lines and two different human primary lung cell culture systems. The dose-proportional pharmacokinetic profile observed after oral administration of GS-621763 translated to dose-dependent antiviral activity in mice infected with SARS-CoV-2. Therapeutic GS-621763 significantly reduced viral load, lung pathology, and improved pulmonary function in COVID-19 mouse model. A direct comparison of GS-621763 with molnupiravir, an oral nucleoside analog antiviral currently in human clinical trial, proved both drugs to be similarly efficacious. These data demonstrate that therapy with oral prodrugs of remdesivir can significantly improve outcomes in SARS-CoV-2 infected mice. Thus, GS-621763 supports the exploration of GS-441524 oral prodrugs for the treatment of COVID-19 in humans.

## Introduction

SARS-CoV-2 emerged in December 2019 and has caused 223 million infections and 4.6 million deaths worldwide as of September 2021 (*1–3*). While there are multiple effective vaccines, vaccination rates have lagged in the United States (U.S.) due to vaccine hesitancy and public mistrust thus delaying the generation of herd immunity required to significantly diminish community spread. In addition, outside of the U.S., many countries do not have equitable access to vaccines and/or have been slow to vaccinate (*4–7*). This collective constellation of events is fueling the generation of viral variants that are increasingly transmissible and are ever evolving to escape human immunity. Therefore, there is an immediate unmet need for oral antivirals that can be rapidly disseminated to treat COVID-19 cases in the unvaccinated, the immunocompromised and in vaccine breakthrough cases. Next-generation oral coronavirus (CoV) antivirals, if widely disseminated and given early in infection, could curtail the duration of disease, reduce long-term sequelae of COVID-19, minimize household transmissions, and lessen hospitalizations, thus having a broad impact on public health.

There are multiple direct-acting antiviral (DAA) therapies in use to treat COVID-19 (*8–12*), including Emergency Use Authorization (EUA)-approved monoclonal antibodies (mAbs) and FDA-approved remdesivir (RDV, GS-5734). Monoclonal antibodies have demonstrated efficacy for treating active COVID-19 cases in outpatients (Regeneron outpatient studies) but currently, all mAbs must be administered via injection, limiting their use to those with ready access to healthcare (*13, 14*). In addition, several SARS-CoV-2 variants of concern (VOCs) have evolved that are resistant to first-line mAb therapies (*15, 16*). Currently, RDV is the only FDA-approved small-molecule direct-acting antiviral to treat COVID-19, but there are several other DAAs currently in human clinical trials including nucleoside analogs molnupiravir (MPV, EIDD-2801) and AT-527, as well as MPro inhibitor PF-07321332 (*17–25*). Unlike mAbs which specifically target the virion surface exposed spike protein of SARS-CoV-2, nucleoside analog drugs target a highly conserved viral enzyme among CoV, the RNA-dependent RNA polymerase (RdRp) nsp12, rendering them broadly active against multiple emerging, endemic and enzootic CoV. Moreover, due to its high degree of conservation among CoV, the RdRp likely does not have the same capacity for mutational change as spike, which may translate into RdRp having a higher barrier to resistance (*18–20*). Despite demonstrated therapeutic efficacy of RDV against SARS-CoV-2 in animal models (*17, 23, 26*) and in human clinical trials (*8*), the requirement of intravenous administration has limited its widespread use during this pandemic. The orally bioavailable nucleoside prodrug GS-621763, is designed for optimal delivery of the parent nucleoside GS-441524 into systemic circulation, which is then metabolized inside cells into the same active nucleoside triphosphate formed by RDV (*27*). Here, we detail the *in vitro* antiviral activity in various cell models and *in vivo* therapeutic efficacy of oral GS-621763 in a mouse model of SARS-CoV-2 pathogenesis.

## Results

### GS-621763 has antiviral activity against SARS-CoV-2 in cell lines and human primary cell cultures

GS-441524 is the parental adenosine nucleoside analog (Fig. 1A) of both monophosphoramidate prodrug RDV (GS-5734, Fig. 1B), and triester prodrug GS-621763 (Fig. 1C). All three molecules are metabolized to the same active nucleotide triphosphate in cells, but through different activation pathways. GS-621763 is rapidly metabolized during oral absorption to GS-441524, then intracellularly converted by cellular kinases to the analog monophosphate metabolite before further metabolism to the active nucleoside triphosphate. In contrast, the intact phosphoramidate prodrug, RDV, is broken down inside cells directly to the same monophosphate metabolite, effectively bypassing the rate-limiting first phosphorylation step of GS-441524 (*27*). To determine if GS-621763 could inhibit replication of SARS-CoV-2 in cellular assays, we first evaluated its antiviral activity against a SARS-CoV-2 reporter virus expressing nanoluciferase (SARS-CoV-2 nLUC) in A549-hACE2 cells stably expressing the human entry receptor angiotensin-converting enzyme 2 (ACE2) (*28*). With GS-621763, we observed a dose-dependent antiviral effect on SARS-CoV-2 nLUC replication with an average half-maximum effective concentration (EC_50_) of 2.8 μM (Fig. 1D, Fig.1H, and Supplementary Figure 1A). In the same assay, we measured EC_50_ values for the control compound RDV of 0.28 μM, similar to those reported previously in these cells and reflective of the enhanced ability of the phosphoramidate prodrug to rapidly and efficiently generate active triphosphate by bypassing the slower initial phosphorylation step (Fig. 1D, Fig. 1H, Supplementary Figure 1A) (*29*). As was observed in other cell systems, the parental nucleoside, GS-441524, was less potent (EC_50_ = 3.3 μM) than RDV in our assay and was similar in potency to GS-621763. This suggests that the tri-isobutyryl esters of GS-621763 are efficiently cleaved in the assay to release GS-441524 (Fig. 1D, Fig. 1H, Supplementary Figure 1A). Importantly, we did not observe any measurable cytotoxicity of any of the inhibitors in A549-hACE2 cells at concentrations up to 10 μM (Figure 1F, Supplementary Figure 1B). Human primary airway epithelial (HAE) cell cultures model the cellular complexity and architecture of the human conducting airway and are often used to determine if drugs are transported and metabolized in the cells targeted by emerging CoV in vivo (*18*). In HAE cells infected with WT SARS-CoV-2 (*28*) and treated with GS-621763, we observed a dose-dependent and significant reduction in infectious virus production as compared to DMSO-vehicle treated cultures (Fig. 1F). In similarly infected, control compound (i.e. RDV or GS-441524) treated cultures, a significant and dose-dependent reduction in viral titers was also observed (Fig. 1F). GS-621763, RDV, and GS-441524 inhibited reporter SARS-CoV-2 expressing Firefly luciferase (SARS-CoV-2 Fluc) replication in normal human bronchial epithelial (NHBE) cultures with EC_50_ values of 0.125, 0.0371, and 2.454 μM, respectively (Fig. 1G, Fig. 1H). All together, these data show that GS-621763 is transported, metabolized and potently antiviral human primary cell systems that model the tissues targeted by SARS-CoV-2 in humans.

**Figure 1.**
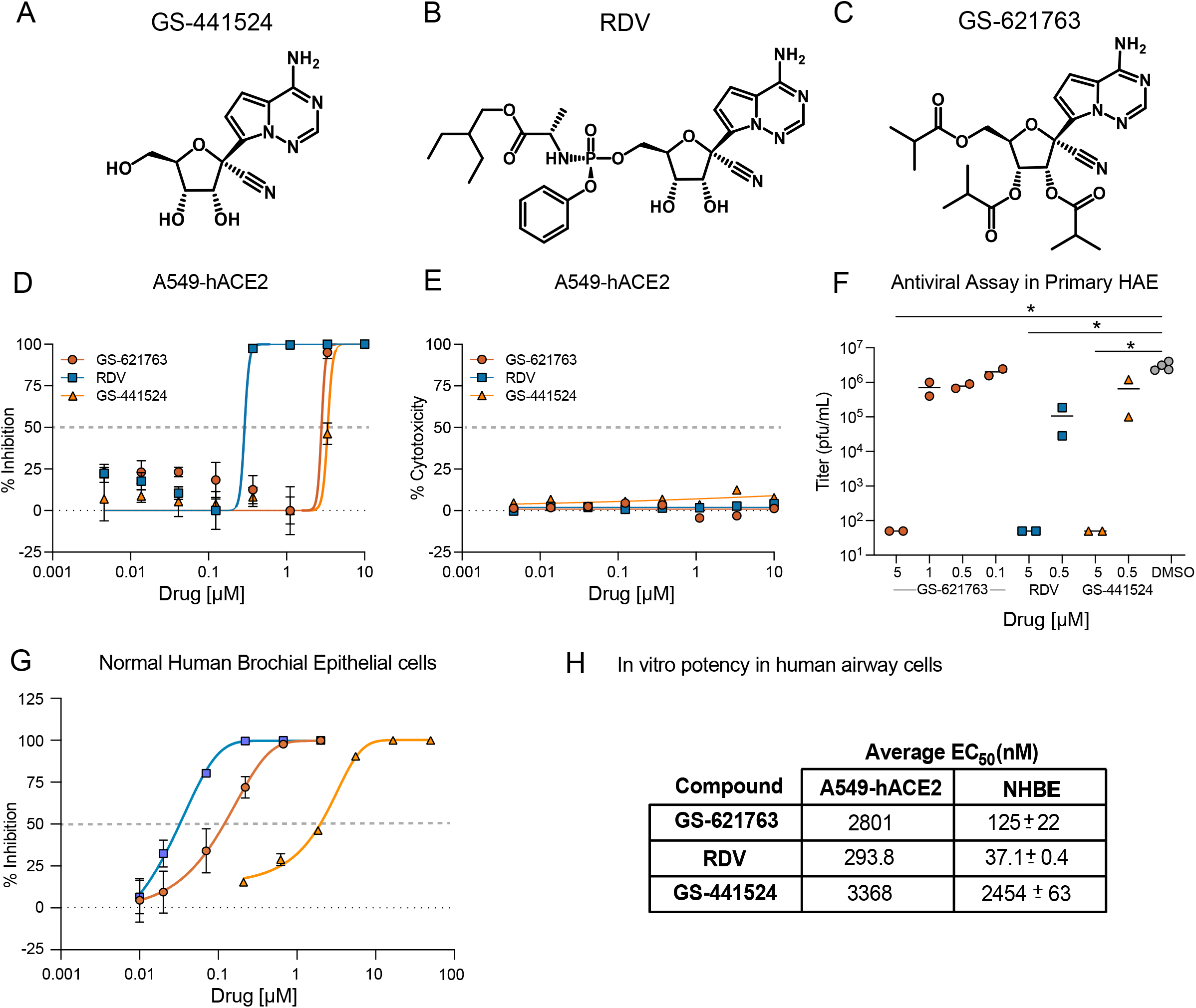
Chemical Structure and in vitro potency of GS-621763 in comparison to RDV (GS-5734) and GS-441524. (A) Chemical structure of the parental adenosine nucleoside analog GS-441524. (B) Chemical structure of the monophosphoramidate prodrug RDV. (C) Chemical structure of GS-621763, the tri-isobutyryl ester of GS-441524. (D) Mean percent inhibition of SARS-CoV-2 replication by GS-621763, in comparison to the prodrug RDV - and the parental nucleoside GS-441524 in A459-hACE2 cells (done in triplicates). (E) Cytotoxicity in A459-hACE2 cells treated with GS-621763, RDV, and GS-441524 in A459-hACE2 cells (done in triplicates, CellTiter-Glo – CTG). (F) Inhibition of SARS-CoV-2 replication by GS-621763, in comparison to the prodrug RDV and the parental nucleoside GS-441524 in human primary airway epithelial cells (HAE, done in duplicates). (G) Inhibition of SARS-CoV-2-Fluc replication by GS-621763, RDV -, and the parental nucleoside GS-441524 in normal human bronchial epithelial (NHBE) cultures (done in duplicates). (H) In vitro EC_50_ values for inhibition of viral replication by GS-621763, RDV, and the parental nucleoside GS-441524 in A459-hACE2 and NHBE cells.

### Dose-dependent therapeutic efficacy of GS-621763 in mouse models of COVID-19 disease

We have previously performed multiple studies describing the therapeutic efficacy of subcutaneously administered RDV in mice (*Ces1c*^−/−^ C57BL/6J) genetically deleted for a secreted plasma carboxylesterase 1c (*Ces1c*) absent in humans but dramatically reduces drug half-life in wild-type mice (*17–19, 26, 30*).

However, the prodrug GS-621763 is designed to be rapidly cleaved pre-systemically in vivo to release GS-441524 into circulation, with no or very minimal intact ester observed in plasma. Therefore, GS-621763 can be studied in wild-type mice where it should also be rapidly converted to parent GS-441524. Plasma pharmacokinetics following a single oral administration of GS-621763 at either 5 or 20 mg/kg were first determined in uninfected BALB/c mice (Fig. 2A). Doses were selected to provide high plasma exposures of GS-441524 that would support active triphosphate formation in the lung and to confirm pharmacokinetic dose proportionality needed to project exposures in efficacy studies. Previous studies had shown that parent nucleoside was at least 10-fold less efficient at generating lung triphosphate than RDV, on a molar basis, thus requiring higher plasma exposures of parental GS-441524 to account for the reduced metabolic efficiency (*27*). No exposure of intact ester prodrug, within the limit of detection, was observed in mice. GS-441524 was both rapidly absorbed and then cleared from systemic circulation, exhibiting a short plasma half-life of approximately 1 hr. Dose proportional increases in both maximal plasma concentrations (C_max_) and exposures (AUC_0-24h_) at the two doses were observed (Fig. 2A).

**Figure 2.**
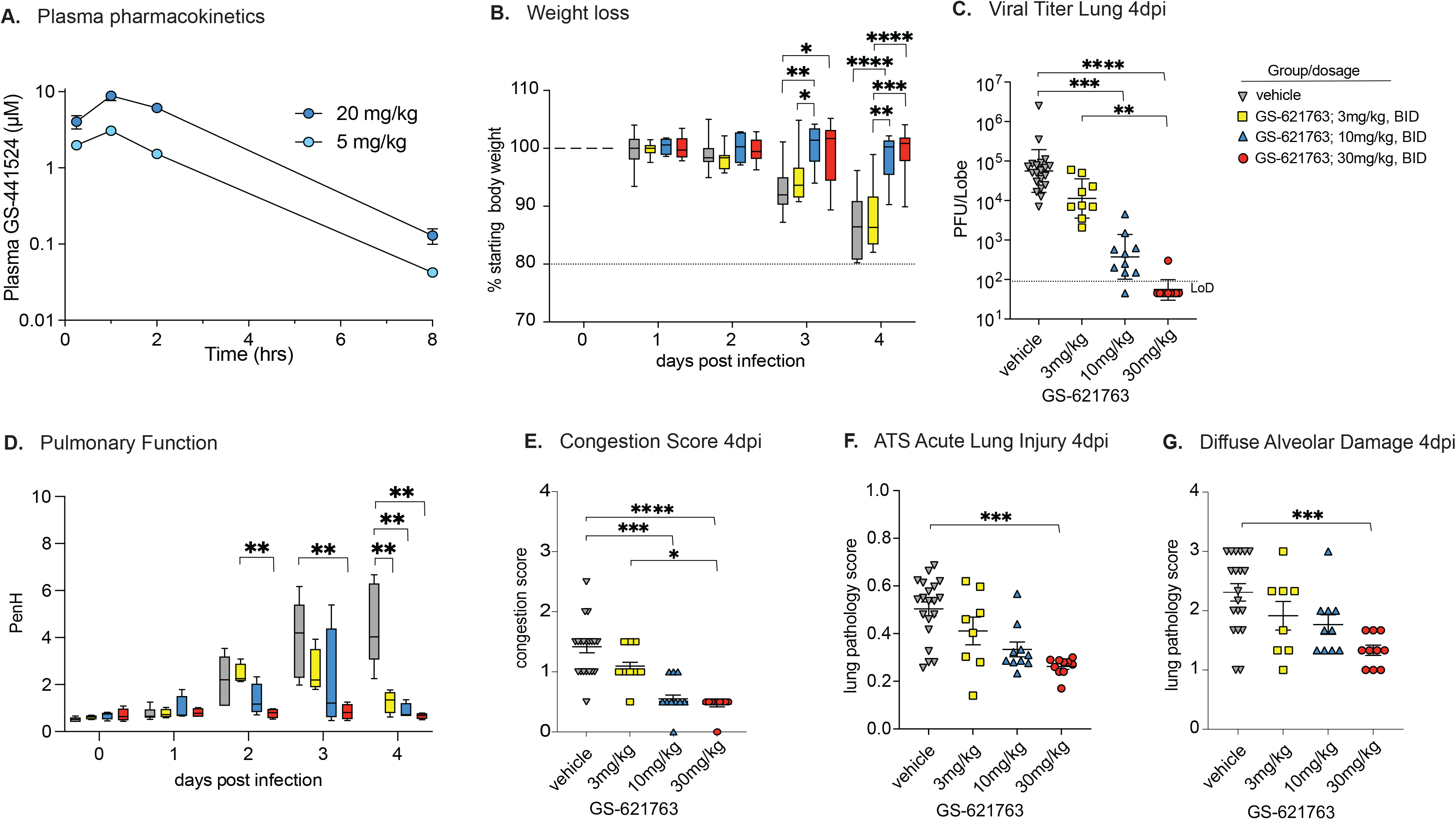
Dose-dependent therapeutic protection against COVID-19 disease by GS-621763 in mice. (A) Plasma Pharmacokinetics of GS-441524 in uninfected Balb/c mice following a single oral administration of GS-621763 at either 5 or 20 mg/kg. Plasma concentrations of GS-621763 were below the limit of quantification at all time points. (B) Depicted is the % starting weight in therapeutically treated mice with vehicle (n=19) or 3 mg/kg (n=9), 10 mg/kg (n=10), and 30 mg/kg (n=10) GS-621763 at 8 hr. All mice were infected with 1×10^4^ PFU SARS-CoV-2 MA10. (C) Lung viral titers in therapeutically treated mice with vehicle (n=19) or 3 mg/kg (n=9), 10 mg/kg (n=10), and 30 mg/kg (n=10) GS-621763 at 8 hr. All mice were infected with 1×10^4^ PFU SARS-CoV-2 MA10. Limit of detection (LoD) (D) Pulmonary function in therapeutically treated mice with vehicle (n=4) or 3 mg/kg (n=4), 10 mg/kg (n=4), and 30 mg/kg (n=4) GS-621763 at 8 hr. All mice were infected with 1×10^4^ PFU SARS-CoV-2 MA10. (E) Lung congestion score in therapeutically treated mice with vehicle (n=19) or 3 mg/kg (n=9), 10 mg/kg (n=10), and 30 mg/kg (n=10) GS-621763 at 8 hr. All mice were infected with 1×10^4^ PFU SARS-CoV-2 MA10. (F and G) Lung pathology in the therapeutically treated mice with vehicle (n=19) or 3 mg/kg (n=9), 10 mg/kg (n=10), and 30 mg/kg (n=10) GS-621763 at 8 hr. All mice were infected with 1×10^4^ PFU SARS-CoV-2 MA10. Data were analyzed using two-way ANOVA (weight loss and lung function) and Kruskal-Wallis test (lung titer, congestion score, and pathology scores), *p<0.05, **p<0.005, ***p<0.0005, ****p<0.0001

To better understand the pharmacokinetic and pharmacodynamic relationship for GS-621763, we performed a series of dose-finding studies in BALB/c mice infected with mouse-adapted SARS-CoV-2 (SARS-CoV-2 MA10) (*30*). In young adult BALB/c mice infected with 10^4^ plaque forming units (PFU) SARS-CoV-2 MA10, virus replicates to high titers in the respiratory tract, mice lose 15-20% of their body weight by 4 days post-infection (dpi), and acute lung injury/loss of pulmonary function is typically observed after virus replication peaks on 2 dpi (*30*). We first defined the minimum dosage sufficient for maximal therapeutic efficacy in BALB/c mice initiating twice daily (i.e. bis in die, BID) oral treatment with either vehicle control or 3 mg/kg, 10 mg/kg, or 30 mg/kg GS-621763 beginning 8 hours post infection (hpi) with 10^4^ PFU SARS-CoV-2 MA10 (Fig. 2B). Unlike vehicle or 3 mg/kg GS-621763 treated animals, mice receiving either 10 or 30 mg/kg GS-621763 were completely protected from weight loss thus demonstrating that early oral antiviral therapy can prevent the progression of disease (Fig. 2B). Congruent with the weight loss phenotype, both 10 and 30 mg/kg GS-621763 treated animals had significantly reduced viral lung titers as compared to both the vehicle and 3 mg/kg treated groups (Fig. 2C). To monitor the effect of drug treatment on pulmonary function, we performed daily whole-body plethysmography (WBP) with a subset of mice from each group (N=4 per treatment group). As shown with the WBP metric PenH, whose elevation is associated with airway resistance or obstruction (*18*), we observed a drug dose-dependent reduction in PenH with the maximal effect seen in the 30 mg/kg GS-621763 dose group which was completely protected from the loss of pulmonary function observed in the other treatment groups and vehicle (Fig. 2D). Mice treated with 3 and 10 mg/kg GS-621763 had impaired lung function at days 2 and 3 post infection, but lung function returned to baseline by 4 dpi for all GS-621763 treated animals (Fig. 2D). Consistent with weight loss, virus titer, and pulmonary function data, mice treated with 10 or 30 mg/kg had significantly reduced lung congestion, a gross pathologic feature characteristic of severe lung damage (Fig. 2E). We then scored lung tissue sections for the histologic features of acute lung injury (ALI) using two complementary semiquantitative tools. First, using an ALI scoring tool created by the American Thoracic Society (ATS), we blindly evaluated three diseased fields per lung section for several features of ALI including alveolar septal thickening, neutrophils in the interstitium and in air spaces, proteinaceous debris in airspaces, and the presence of hyaline membranes. Only mice treated with 30 mg/kg had significantly reduced ALI scores (Fig. 2F). Second, we used a complementary tool measuring the pathologic hallmark of ALI, diffuse alveolar damage (DAD). Mice in all treated groups showed reduced DAD scores, but only mice receiving 30 mg/kg had significantly decreased DAD in their lungs (Fig. 2G). Together, these data demonstrate that the oral delivery of the nucleoside analog GS-621763 can significantly diminish SARS-CoV-2 virus replication and associated pulmonary disease in a dose-dependent manner.

### Extended therapeutic protection against COVID-19 disease by GS-621763 in mice

To determine if the potent therapeutic efficacy of GS-621763 observed with early intervention (8 hr after infection) would extend to later times post infection, we designed a therapeutic efficacy study with six arms where we varied both time of oral therapy initiation and dose level in BALB/c mice infected with SARS-CoV-2 MA10 (Fig.3). As done previously, a control group of animals received vehicle twice daily beginning at 12 hours post infection (hpi). The next three arms of the study were dedicated to the 30 mg/kg GS-621763 dose level, with two of the three arms receiving twice daily dosing initiated at either the 12 hpi (“30 mg/kg BID 12 hr” group) or the 24 hpi (“30 mg/kg BID 24 hr” group). The third 30 mg/kg arm was designed to determine if dose frequency could be reduced to once daily (quaque die, QD) if initiated early at 12 hpi (“30 mg/kg QD 12 hr” group). In the last two arms, we wanted to evaluate if an increased dose of 60 mg/kg given QD beginning at 12 hr or 24 hr (“60 mg/kg QD 12 hr” and “60 mg/kg QD 24 hr” groups) would improve outcomes over the 30 mg/kg groups. Initiation of 30 mg/kg BID therapy at either 12 or 24 hrs offered significant protection from weight loss (Fig. 3A), extending the robust therapeutic phenotype observed for this dose level when initiated at very early times (at 8 hr) (Fig 2). Interestingly, when we decreased the frequency of 30 mg/kg treatment initiated at 12 hr to once daily (“30 mg/kg QD 12 hr” group), we also observed a significant prevention of body weight loss (Fig 3A), thus levels of drug when administered once a day and begun early (at 12 hr) in the course of infection were sufficient to prevent disease progression. Increasing the dose to 60 mg/kg QD initiated at either 12 hr or 24 hr offered similar protection from weight loss observed with vehicle treatment as the 30 mg/kg groups (Fig. 3A).

**Figure 3.**
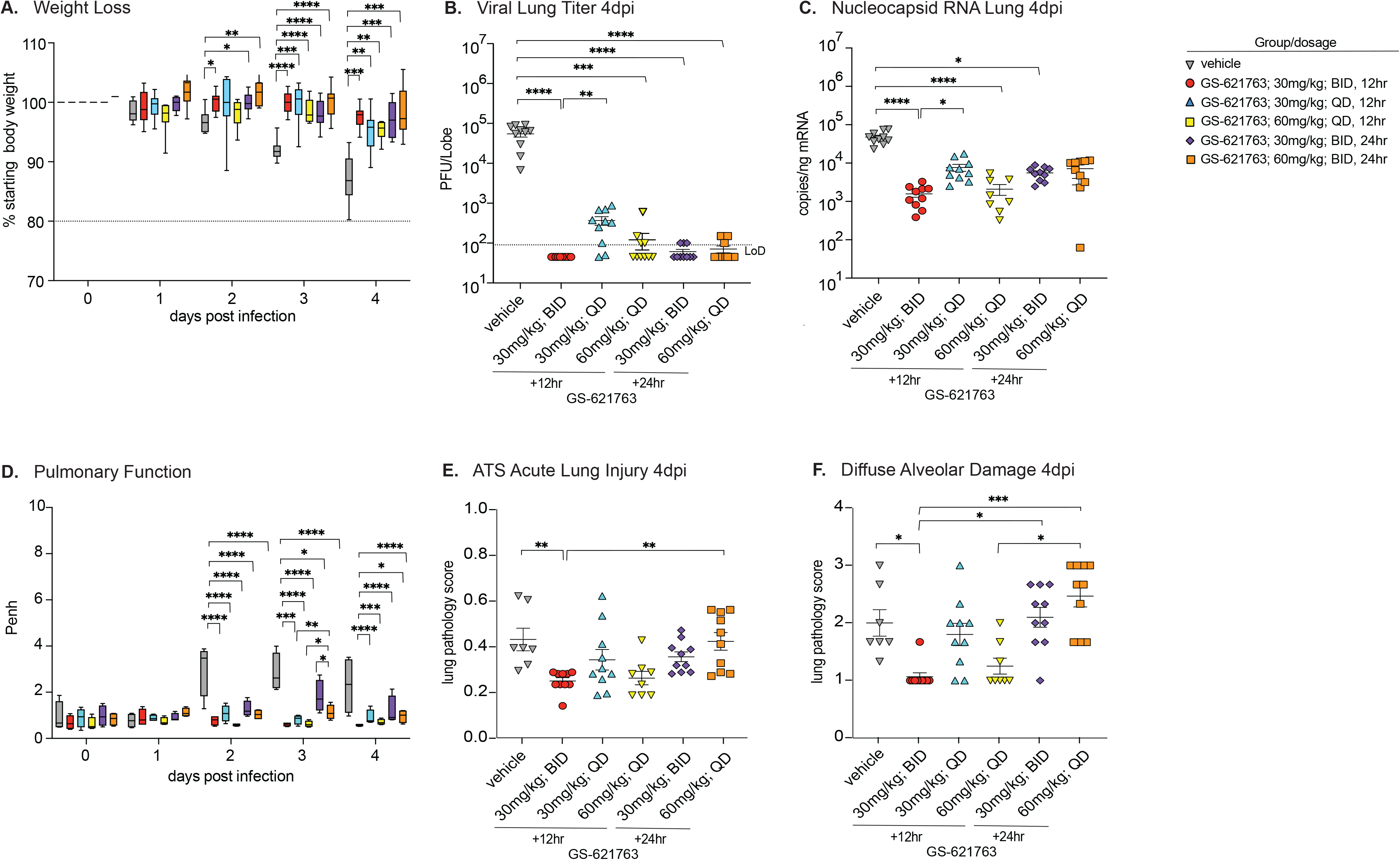
Extended therapeutic protection of mice against COVID-19 disease by oral GS-621763 in mice. (A) Depicted is the % starting weight in therapeutically treated mice with either 30 mg/kg or 60 mg/kg BID or QD GS-621763 at 12 and 24 hrs (n=10 for all treatment groups, except n=8 for 60 mg/kg, 12 hr QD treatment group). All mice were infected with 1×10^4^ PFU SARS-CoV-2 MA10. (B) Lung viral titers in therapeutically treated mice with either 30 mg/kg or 60 mg/kg BID or QD GS-621763 at 12 and 24hrs (n=10 for all treatment groups, except n=8 for 60 mg/kg, 12 hr QD treatment group). All mice were infected with 1×10^4^ PFU SARS-CoV-2 MA10. Limit of detection (LoD). (C) Viral N RNA in therapeutically treated mice with either 30 mg/kg or 60 mg/kg BID or QD GS-621763 at 12 and 24 hrs (n=10 for all treatment groups, except n=8 for 60 mg/kg, 12 hr QD treatment group). All mice were infected with 1×10^4^ PFU SARS-CoV-2 MA10. (D) Pulmonary function in therapeutically treated mice with either 30 mg/kg or 60 mg/kg BID or QD GS-621763 at 12 and 24 hrs (n=4 for all treatment groups). All mice were infected with 1×10^4^ PFU SARS-CoV-2 MA10. (n=4 for all treatment groups). (F and G) Lung pathology in the therapeutically treated mice with either 30 mg/kg or 60 mg/kg BID or QD GS-621763 at 12 and 24 hrs (n=10 for all treatment groups, except n=8 for 60 mg/kg, 12 hr QD treatment group, and n=7 for vehicle group). Data were analyzed using two-way ANOVA (weight loss and lung function) and Kruskal-Wallis test (lung titer, lung viral RNA, and pathology scores), *p<0.05, **p<0.005, ***p<0.0005, ****p<0.0001.

As body weight loss is a crude marker of viral pathogenesis, we next measured multiple virological, physiologic, and pathologic metrics of disease. First, we measured the levels of infectious virus present in lung tissue on 4 dpi. Unlike vehicle-treated animals who harbored an average titer of 5.6×10^4^ PFU per lung lobe, all GS-621763 dose groups significantly reduced the levels of infectious virus in lung tissue with the average titers of most groups falling below the limit of detection (90 PFU). Interestingly, 30 mg/kg delivered QD had significantly elevated viral lung titers (mean titer = 4×10^2^ PFU) as compared to its BID counterpart (mean titer = < 90 PFU). A similar trend among treatment groups was observed when measuring the levels of SARS-CoV-2 subgenomic and genomic nucleocapsid (N) RNA in parallel lung tissues (Fig. 3C). All 30 mg/kg groups significantly reduced levels of SARS-CoV-2 RNA in lung tissue as compared to vehicle-treated animals. As observed for infectious titers, 30 mg/kg given QD daily beginning at 12 hr had elevated levels of N RNA as compared to 30 mg/kg given BID beginning at 12 hr suggesting that trough and/or daily exposure levels of drug with QD dosing are insufficient to suppress replication similarly to BID dosing. Increasing the once-daily dose to 60 mg/kg to raise the daily exposure and trough levels offered similar reductions in SARS-CoV-2 N RNA as compared to 30 mg/kg BID when initiated at 12 hr, but the levels of viral RNA in the higher dose 60mg/kg group initiated at 24 hr were not different than vehicle (Fig. 3C). Although vehicle-treated animals exhibited significant loss of pulmonary function as measured by WBP, this was largely prevented with GS-621763 therapy (Fig. 3D). All therapy groups initiated at 12 hr were equally protected from loss of pulmonary function as measured by the PenH metric (Fig. 3D). Of all groups actively receiving GS-621763, animals in the 30 mg/kg BID 24 hr group had a measurable loss of lung function on 3 dpi which resolved by 4 dpi but this phenotype was not extended to other groups. We then scored lung tissue sections for the histologic features of acute lung injury and alveolar damage. Only 30 mg/kg BID initiated at 12 hr significantly reduced ALI scores as compared to those in vehicle-treated animals. In addition, the 30 mg/kg BID 12 hr-group had significantly lower ALI scores as compared to the 60 mg/kg QD 24 hr-group (Figure 3E). Only 30 mg/kg given twice a day at 12 hr most dramatically reduced DAD scores but this protection from lung pathology was lost if given once per day or if initiated at 24 hr (Fig. 3G). The high dose of 60 mg/kg QD when initiated at 12 hr improved DAD scores as compared to similarly treated animals that began treatment at 24 hr. Collectively, these data demonstrate that GS-621763 therapy can improve both virologic and pathogenic metrics, but the degree of improvement was dependent on time of initiation and dose frequency.

### The therapeutic efficacy of GS-621763 is similar to molnupiravir (MPV, EIDD-2801)

MPV is an oral nucleoside analog prodrug antiviral currently in Phase 3 clinical trial to treat COVID-19 with demonstrated antiviral efficacy in mice against several emerging CoV including SARS-CoV, MERS-CoV, and SARS-CoV-2 (*18, 19, 21, 22*). Like GS-621763, MPV is a prodrug which is metabolized in vivo into a parental nucleoside (β-D-N4-hydroxycytidine, NHC) in its metabolic progression towards the antiviral active triphosphate (*20*). To determine if GS-621763 would provide similar protection as MPV, we then designed comparative therapeutic efficacy studies in the mouse model of SARS-CoV-2 pathogenesis described above. Pre-efficacy pharmacokinetic studies in BALB/c mice (30 mg/kg or 100 mg/kg) were performed with MPV and showed dose proportional increases in NHC plasma exposures (Supplemental Fig. 2). Pharmacokinetic modeling then determined that a daily 120 mg/kg dose (given 60 mg/kg BID) would result in exposures similar to that observed in humans receiving 800 mg BID, a dose being evaluated in a human clinical trial (*22*). The comparative efficacy study included a vehicle group and 5 additional groups receiving two doses of MPV or GS-621763 per day 12 hrs apart (BID). Three arms of the study began dosing at 12 hr: 30 mg/kg GS-621763, 30 mg/kg MPV (0.5× human equivalent dose) or 60 mg/kg MPV (1× human equivalent dose). At 24 hr, we began dosing of two additional groups: 60 mg/kg GS-621763 or 60 mg/kg MPV. While SARS-CoV-2 MA10 infection caused rapid weight loss in vehicle control animals, all animals receiving either GS-621763 or MPV beginning at either 12 or 24 hr were protected from weight loss (Fig. 4A). Similarly, upon titration of lung tissues at 4 dpi for infectious virus by plaque assay, vehicle-treated animals had expectedly high levels of infectious virus which was significantly reduced in all treatment groups, independent of drug type or initiation time (Fig. 4B). When treatment was initiated at 12 hr, a moderate yet significant elevation in infectious titers was observed in the 30mg/kg MPV group, inferior to either equivalently dosed GS-621763 animals or those receiving the higher dose (60 mg/kg) of MPV (Fig. 4B). To understand the relationship between levels of infectious virus and viral RNA in lung tissue, we performed qRT-PCR on total RNA for SARS-CoV2 N RNA in parallel tissues utilized for plaque assay. The trend observed with infectious virus is mirrored in the qRT-PCR data where all groups receiving antiviral therapy had significantly reduced levels of viral RNA (Fig. 4C). In addition, animals receiving 30 mg/kg MPV (0.5× human equivalent dose) had a measurable increase in N RNA as compared to equivalently dosed GS-621763 animals. Similar to weight loss data, vehicle-treated animals had a significant loss of pulmonary function as measured by WBP on both 3 and 4 dpi which was prevented in all groups receiving antiviral treatment (Fig. 4D). We then blindly evaluated lung tissue sections for the pathological manifestations of ALI and DAD using two complementary histologic tools described above. Congruent with the above data, ALI scores in all antiviral therapy groups were significantly reduced as compared to vehicle controls (Fig. 4E). In agreement with ALI scores, the DAD histologic scores were similarly reduced in all antiviral therapy treated groups as compared to those treated with vehicle (Fig. 4F). All together, these data show that antiviral therapy with GS-621763 and MPV when initiated early or at the peak of virus replication (~24 hr) can both significantly diminish virus replication and improve disease outcomes.

**Figure 4.**
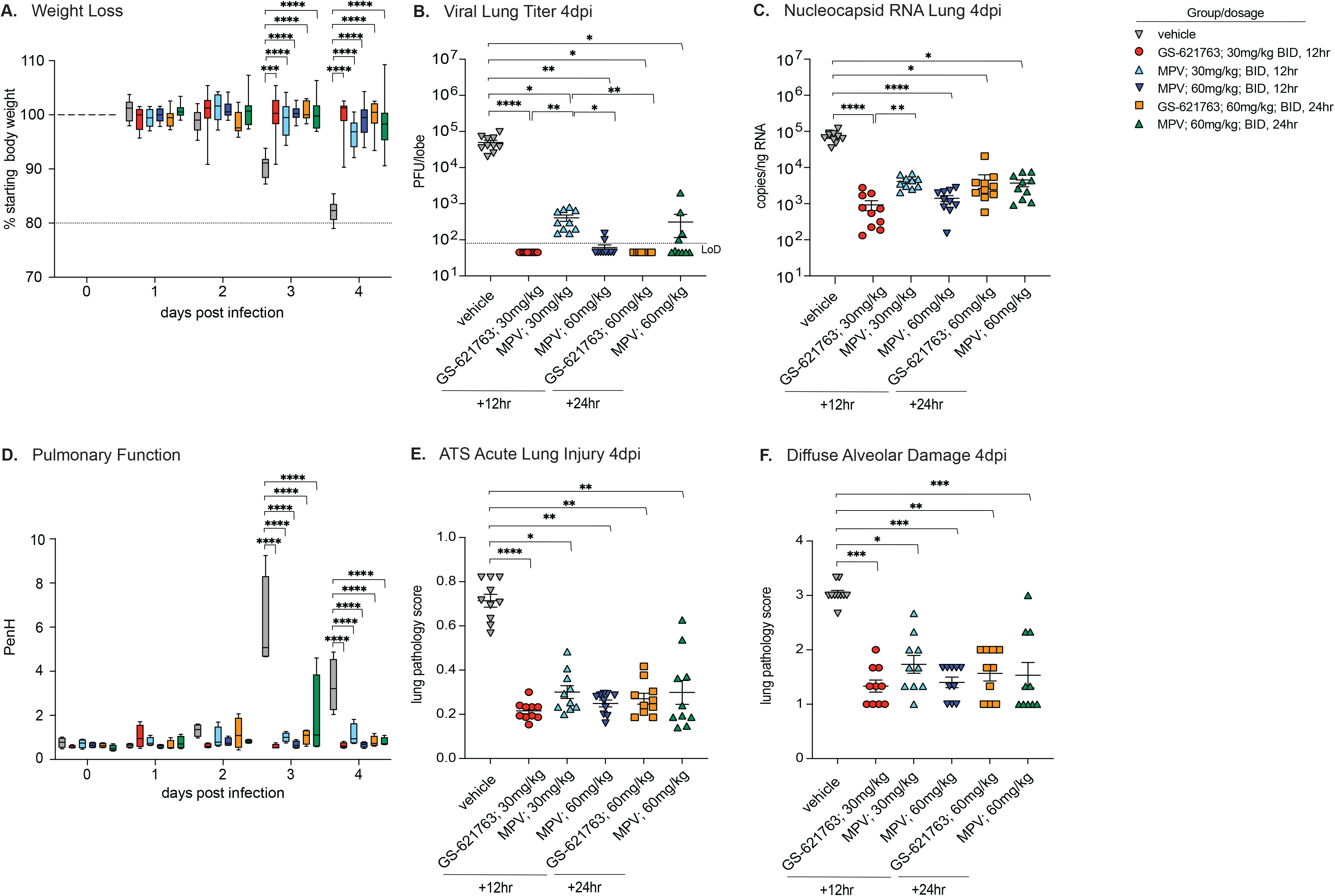
Evaluation of therapeutic intervention of GS-621763 in comparison to molnupiravir (MPV). (A) Depicted is the % starting weight in therapeutically treated mice with either 30 mg/kg or 60 mg/kg GS-621763 or MPV at 12 or 24 hrs (n=10 for all treatment groups). All mice were infected with 1×10^4^ PFU SARS-CoV-2 MA10. (B) Lung viral titers in therapeutically treated mice with either 30 mg/kg or 60 mg/kg GS-621763 or MPV at 12 or 24 hrs (n=10 for all treatment groups). All mice were infected with 1×10^4^ PFU SARS-CoV-2 MA10. Limit of detection (LoD). (C) Viral N RNA in therapeutically treated mice with either 30 mg/kg or 60 mg/kg GS-621763 or MPV at 12 or 24 hrs (n=10 for all treatment groups). All mice were infected with 1×10^4^ PFU SARS-CoV-2 MA10. (D) Pulmonary function in therapeutically treated mice with either 30 mg/kg or 60mg/kg GS-621763 or MPV at 12 or 24 hrs (n=4 for all treatment groups). All mice were infected with 1×10^4^ PFU SARS-CoV-2 MA10. (F and G) Lung pathology in the therapeutically treated mice with either 30 mg/kg or 60 mg/kg GS-621763 or MPV at 12or 24 hrs (n=10 for all treatment groups). All mice were infected with 1×10^4^ PFU SARS-CoV-2 MA10. Data were analyzed using two-way ANOVA (weight loss and lung function) and Kruskal-Wallis test (lung titer, lung viral RNA, and pathology scores), *p<0.05, **p<0.005, ***p<0.0005, ****p<0.0001

## Discussion

Three novel human CoVs have emerged in the past 20 years, first with SARS-CoV in 2002-2003, MERS-CoV in 2012 and most recently, SARS-CoV-2 in 2019 (*1, 2, 31*). Vaccine availability, vaccine hesitancy, and the ongoing evolution and emergence of VOCs are collectively delaying the global control of pandemics and potential achievement of global herd immunity and may prevent it altogether (*15, 32–34*). As such, there is an acute need for broad-spectrum antivirals to treat COVID-19 in the unvaccinated as well as increasingly common breakthrough infections in those vaccinated, driven by immune evading VOCs. In addition, due to the emergence potential of the CoV family, we must also actively develop broadly acting therapies effective the CoVs of today to prepare for those that may emerge in the future. Both RDV and MPV are examples of nucleoside analogs with broad activity against genetically diverse CoV (*35*). Here, we show that GS-621763, a prodrug that improves the oral delivery of parent nucleoside GS-441524, is yet another example of an orally bioavailable nucleoside analog prodrug that is effective against SARS-CoV-2.

Oral broadly acting antiviral therapies that target conserved viral proteins with a diminished capacity for evolutionary change will maximize therapeutic utility against future emerging CoV. SARS-CoV-2 has undergone a considerable amount of genetic evolution since its emergence in 2019; each time a globally dominant SARS-CoV-2 VOC has emerged, it has been replaced by a new VOC harboring more concerning characteristics like replicative capacity or transmissibility (*36*). The majority of genetic changes have been localized to viral proteins decorating the surface of the virus particle like the viral spike which has mutated to eradicate epitopes targeted by early and promising monoclonal antiviral therapies (e.g. Eli Lily, Regeneron) rendering them less active against newer VOCs (*37*). Approved antivirals like RDV, those in clinical trials like MPV, AT-527, and PF-07321332 (Pfizer, oral protease inhibitor) target highly conserved enzymes required for virus replication which likely have a diminished capacity for change as compared to the spike protein (*38*). The widespread use of both RDV and mAb therapies in the U.S. and globally has been limited by the necessity of delivery by intravenous infusion which in turn requires access to qualified health care staff and facilities. Thus, effective oral antiviral therapy or combination therapies which can be procured at a pharmacy and self-administered by the patient would help facilitate wide-spread global access and could have profound positive impacts on global public health.

In this study, we utilized a mouse adapted SARS-CoV-2 variant, SARS-CoV-2 MA10, in wild-type BALB/c mice. (*17, 30*). Mice infected with this virus develop severe lung disease reminiscent of that seen with severe COVID-19 including the development of ALI and respiratory failure (*30*). It is important to note that the disease resulting from SARS-CoV-2 MA10 infection in mice is compressed as compared to that observed in humans with virus titer peaking in the mouse lung between 24-48 hrs after infection, rapid loss of pulmonary function beginning 2-3 dpi, rapid weight loss within the first 4 days of infection, lung pathology consistent with ALI peaking 4-6 dpi and virus induced mortality within a week of infection. This is markedly different than the COVID-19 in humans where virus titers peak in the upper airway within the first week, but viral RNA shedding can be observed for as long as 24 days and symptoms can take weeks to months to resolve (*39–41*). Because of this caveat associated with our mouse model, the time in which to intervene with a direct acting antiviral and sufficiently improve outcomes is curtailed in mice as compared to humans. Our recent study with RDV exemplifies this where we found the degree of therapeutic benefit in mice infected with SARS-CoV-2 MA10 was dependent on the time of initiation (*17*). Here, we show that oral administration with GS-621763 prevents body weight loss, loss of pulmonary function, severe lung pathology and virus replication when administered at 12 and 24 hrs after infection. While we observed improvement in some metrics with RDV initiated at 24 hr in our prior studies (*17*), GS-621763 therapy initiated at a similar time comparatively improved all metrics assessed. Thus, herein we provide proof-of-concept preclinical data that the orally bioavailable ester analog of RDV GS-621763 can exert potent antiviral effects in vivo during an ongoing SARS-CoV-2 infection. Lastly, we show that GS-621763 therapy provides similar levels of protection from SARS-CoV-2 pathogenesis as MPV, an oral nucleoside analog prodrug effective against SARS-CoV-2 in mice that is currently in human clinical trials. Future directions are focused on extending these studies to evaluate the efficacy of combinations of antivirals in our models of SARS-CoV-2 pathogenesis and in other models that can evaluate the blockade of transmission such as hamster and ferret.

In summary, we provide preclinical data demonstrating the in vitro antiviral activity and in vivo therapeutic efficacy of an orally bioavailable nucleoside analog prodrug, GS-621763. The data provided herein supports the future evaluation of orally bioavailable prodrugs of GS-441524 in humans with COVID-19. If safe and effective, this class of RdRp inhibitors could become part of the arsenal of existing oral antivirals that are desperately needed to address a global unmet need for the COVID-19 pandemic and CoV pandemics of the future.

## Figure Legends

**Supplemental Figure 1. In vitro potency and toxicity of GS-621763, RDV, and GS-441524 in A549-hACE2 cells**

(A) Raw data for the inhibition of SARS-CoV-2 replication by GS-621763, RDV, and GS-441521 in A459-hACE2 cells measured through quantitation of SARS-CoV-2 expressed nano luciferase (nLuc), measured in triplicates.

(B) Raw data for cytotoxicity in A459-hACE2 cells treated with GS-621763, RDV, and GS-441521 in A459-hACE2 cells measured via CellTiter-Glo, (measured in triplicates).

C) Raw data for the inhibition of SARS-CoV-2-Fluc replication by GS-621763, RDV, and GS-441521 in NHBE cultures measured through quantitation of SARS-CoV-2 expressed firefly luciferase (Fluc), measured in duplicates, repeated twice.

**Supplemental Figure 2. Molnupiravir mouse plasma pharmacokinetics**

Plasma Pharmacokinetics of N-hydroxycytidine (NHC) in uninfected Balb/c mice following daily oral administration of molnupiravir at either 60 or 200 mg/kg (as either 30 or 100 mg/kg/dose given BID; molnupiravir at all timepoints).

## Material and Methods

### Small molecule drug synthesis and formulation

GS-621763, RDV, and GS-441524 were synthesized at Gilead Sciences Inc., and their chemical identity and purity were determined by nuclear magnetic resonance, and high-performance liquid chromatography (HPLC) analysis (*27*). Molnupiravir was purchased from MedChemExpress LLC (NJ, USA) with a purity of 95% based on HPLC analysis.

Small molecules were solubilized in 100% DMSO for in vitro studies and in vehicle containing 2.5% DMSO; 10% Kolliphor HS-15; 10% Labrasol; 2.5% Propylene glycol; 75% Water (final formulation pH 2) (for GS-621763) and in vehicle containing 2.5% Kolliphor RH-40, 10% Polyethylene glycol 300, 87.5% Water (for MPV) for in vivo studies. GS-621763, GS-441524, GS-5734 were made available to the University of North Carolina (UNC) at Chapel Hill under a materials transfer agreement with Gilead Sciences.

### In vivo plasma pharmacokinetic analysis of GS-621763 and molnupiravir (MPV)

Mice were orally administered either a single dose of GS-621763 (in vehicle containing 2.5% DMSO; 10% Kolliphor HS-15; 10% Labrasol; 2.5% Propylene glycol; 75% Water (final formulation pH 2) or two doses of molnupiravir (in vehicle containing 2.5% Kolliphor RH-40, 10% Polyethylene glycol 300, 87.5% Water) (BID, 12 hours apart). GS-621763 was given at either 5 or 20 mg/kg and MPV at either 30 or 100 mg/kg. Plasma was serially isolated from 4 mice at 0.25, 1, 2, 8 and 24 hrs post GS-621763 administration. Plasma was isolated from alternating groups of 4 mice per timepoint at 0.5, 2, 6, 12 (pre-second dose), 12.5, 18 and 24 hrs post MPV administration. 20 μl of plasma was added to a mixture containing 250 μl of methanol and 25 μL of internal standard solution and centrifuged. 250 μl of resulting supernatant was then transferred, filtered (Agilent Captiva 96, 0.2 μm) and dried under a stream of nitrogen at 40 oC. Following reconstitution in a mixture of 5% acetonitrile and 95% water, a 10 μl aliquot was injected onto an LC-MS/MS system. Plasma concentrations of either GS-621763 and GS-441524 or MPV and N-hydroxycytidine (NHC) were determined using 8 to 10-point calibration curves spanning at least 3 orders of magnitude with quality control samples to ensure accuracy and precision, prepared in normal mouse plasma. Analytes were separated by a 50 mm × 3.0 mm, 2.55 μm Synergi Polar-RP column (Phenomenex) using a multi-stage linear gradient from 5% to 95% acetonitrile in mobile phase A at a flow rate of 1 ml/min.

### Quantitation of GS-441524 metabolites in the lung following oral GS-621763 administration in *Balb/c* mice

Lungs from all mice administered GS-621763 were quickly isolated at 24 hrs post-dose and immediately snap frozen in liquid nitrogen. On dry ice, frozen lung samples were pulverized and weighed. Dry ice-cold extraction buffer containing 0.1% potassium hydroxide and 67 mM ethylenediamine tetraacetic acid (EDTA) in 70% methanol, containing 0.5μM chloro-adenosine triphosphate as internal standard was added and homogenized. After centrifugation at 20,000 × g for 20 minutes, supernatants were transferred and dried in a centrifuging evaporator. Dried samples were then reconstituted with 60 μL of mobile phase A, containing 3 mM ammonium formate (pH 5) with 10 mM dimethylhexylamine (DMH) in water, centrifuged at 20,000 × g for 20 minutes and final supernatants transferred to HPLC injection vials. An aliquot of 10 μl was subsequently injected onto an API 6500 LC/MS/MS system for analysis of GS-441524 and its phosphorylated metabolites, performed using a similar method as described previously (*18*).

### Viruses and plaque assay

Recombinant SARS-CoV-2 MA10 virus was generated as described previously (*30*). For virus titration by plaque assay, the caudal lobe of the right lung was homogenized in PBS, and the resulting homogenate was serial-diluted and inoculated onto confluent monolayers of Vero E6 cells, followed by agarose overlay. Plaques were visualized with overlay of neutral red dye on day 3 after infection (*30*).

### In vitro assays for antiviral activity

A549-hACE2 cells were plated at a density of 20,000 cells/well/100 μl in black-walled clear-bottom 96-well plates 24 hrs prior to infection. Compounds GS-621763, GS-5734, GS-441524, were diluted in 100% DMSO (1:3) resulting in a 1000X dose response from 10 to 0.002 mM (10 to 0.002 μM final). All conditions were performed in triplicate. At BSL3, medium was removed, and cells were infected with 100 μl SARS-CoV-2 nLUC (MOI 0.008) for 1 h at 37 °C after which virus was removed, wells were washed (150 μl) with infection media (DMEM, 4% FBS, 1X antibiotic/antimycotic) and infection media (100 μl) containing a dose response of drug was added. Plates were incubated at 37 °C for 48 hrs. NanoGlo assay was performed 48 hpi. Sister plates were exposed to drug but not infected to gauge cytotoxicity via CellTiter-Glo assay (CTG, Promega, Madison, WI), 48 hrs post treatment.

Normal human bronchial epithelial (NHBE) cells (donor 41219) were purchased from Lonza (Walkersville, MD Cat# CC-2540) and maintained in Bronchial Epithelial Cell Growth Medium (BEGM) (Lonza, Walkersville, MD, Cat# CC-3170) with all provided supplements in the BulletKit. Cells are passaged 2-3 times per week to maintain sub-confluent densities and are used for experiments at passages 2-4. NHBE cells were seeded in 24-well plates at 1×10^5^ cells in a final volume of 0.5 ml BEGM™ Bronchial Epithelial Cell Growth Medium (BulletKit™; Lonza, Basel, SW). Cultures were incubated overnight 37 °C with 5% CO_2_. On the following day, media was replaced with 0.5 ml growth medium. Cultures were treated with 1:3 serial dilutions of compound using the HP D300e digital dispenser with normalization to the highest concentration of DMSO in all wells (<1% final volume). The cells were then infected with 0.1 ml SARS-CoV-2-Fluc diluted in BEGM media at MOI = 5. Uninfected and untreated wells were included as controls to determine compound efficacy against SARS-CoV-2-Fluc. Following incubation with compound and virus for 24 hrs at 37 °C with 5% CO_2_, culture supernatant was removed from each well and replaced with 0.3 ml of ONE-Glo luciferase reagent (Promega, Madison, WI). The plates were shaken at 400 rpm for 10 min at room temperature. 0.2 ml of supernatant from each well was transferred to a 96-well opaque plate (Corning) and luminescence signal was measured using an EnVision plate reader (PerkinElmer). Values were normalized to the uninfected and infected DMSO controls (0% and 100% infection, respectively). Data was fit using a four-parameter non-linear regression analysis using Graphpad Prism. EC50 values were then determined as the concentration reducing the firefly luciferase signal by 50%. The compiled data was generated based on least two independent experimental replicates, each containing four technical replicates for each concentration.

### Mouse studies and in vivo infections

All mouse studies were performed at the University of North Carolina (Animal Welfare Assurance #A3410-01) using protocols (#20-059) approved by the University of North Carolina Institutional Animal Care and Use Committee. All animal work was approved by the Institutional Animal Care and Use Committee at University of North Carolina at Chapel Hill according to guidelines outlined by the Association for the Assessment and Accreditation of Laboratory Animal Care and the US Department of Agriculture. All work was performed with approved standard operating procedures and safety conditions for SARS-CoV-2. Our institutional BSL3 facilities are designed to conform to the safety requirements recommended by Biosafety in Microbiological and Biomedical Laboratories, the US Department of Health and Human Services, the Public Health Service, the Centers for Disease Control and Prevention, and the National Institutes of Health. Laboratory safety plans have been submitted, and the facility has been approved for use by the University of North Carolina Department of Environmental Health and Safety and the Centers for Disease Control and Prevention.

Even groups (n=10, or less as indicated) of 10-week-old female BALB/c mice (Envigo; #047) were used in all in vivo efficacy studies. For infection, mice were anesthetized with a mixture of ketamine/xylazine and infected with 10^4^ PFU of SARS-CoV-2 MA10 in 50 μl PBS intranasally. Vehicle or GS-621673 was administered orally at the dosages and timepoints as indicated. Mice were monitored daily for body weight changes and for lung function by whole-body plethysmography. At 4 dpi, mice were euthanized, and lung tissue was harvested for viral titer analysis, RNA and histology, and lung congestion scores were estimated (*30*). Samples for viral load determination and for RNA isolation were stored at −80 °C until used; histology samples were inactivated in 10% NBF and stored at 4 °C until further processing.

### Histology and lung pathology scoring

Two separate lung pathology scoring scales, Matute-Bello and Diffuse Alveolar Damage (DAD), were used to quantify acute lung injury (ALI) (*19*).

For Matute-Bello scoring samples were blinded and three random fields of lung tissue were chosen and scored for the following: (A) neutrophils in alveolar space (none = 0, 1–5 cells = 1, > 5 cells = 2), (B) neutrophils in interstitial space (none = 0, 1–5 cells = 1, > 5 cells = 2), (C) hyaline membranes (none = 0, one membrane = 1, > 1 membrane = 2), (D) Proteinaceous debris in air spaces (none = 0, one instance = 1, > 1 instance = 2), (E) alveolar septal thickening (< 2Å~ mock thickness = 0, 2–4Å~ mock thickness = 1, > 4Å~ mock thickness = 2). Scores from A–E were put into the following formula score = [(20x A) + (14 x B) + (7 x C) + (7 x D) + (2 x E)]/100 to obtain a lung injury score per field and then averaged for the final score for that sample.

In a similar way, for DAD scoring, three random fields of lung tissue were scored for the in a blinded manner for: 1= absence of cellular sloughing and necrosis, 2= uncommon solitary cell sloughing and necrosis (1–2 foci/field), 3=multifocal (3+foci) cellular sloughing and necrosis with uncommon septal wall hyalinization, or 4=multifocal (>75% of field) cellular sloughing and necrosis with common and/or prominent hyaline membranes. To obtain the final DAD score per mouse, the scores for the three fields per mouse were averaged.

### RNA isolation and RT-qPCR

Mouse tissue from SARS-CoV-2 infected mice was homogenized using glass beads in TRIzol Reagent (Invitrogen). Equal volume of 100% EtOH was mixed with the TRIzol homogenate and processed using Direct-Zol RNA MiniPrep Kit (Zymo) to extract viral RNA. Optional DNase I treatment was conducted to ensure the adequate removal of unwanted DNA. Eluted RNA was coupled with TaqMan Fast Virus 1-step Master Mix (Applied Biosystems) and nCOV_N1 primers/probe (IDT) to quantify viral load via reverse transcription – quantitative polymerase chain reaction (RT-qPCR). Samples were plated on a MicroAmp EnduraPlate (Applied Biosystems) and run using a QuantStudio 6 Real-Time PCR System (Applied Biosystems) to obtain viral titers. The following PCR program was run: 50 °C for 5 minutes, 95 °C for 20 seconds, followed by 45 cycles of 95 °C for 3 seconds and 60°C for 30 seconds. The sequences of the 2019-nCOV_N1 primers and probe were as follows: Forward primer: GAC CCC AAA ATC AGC GAA AT, Reverse primer: TCT GGT TAC TGC CAG TTG AAT CTG, Probe: AC CCC GCA TT ACG TTT GGT GGA CC (CDC N1 qRT-PCR assay (*42*). SARS-CoV-2 standard curve RNA was produced by PCR amplification of SARS-CoV-2 nucleocapsid by which a 5’ T7 polymerase promoter was introduced. This amplicon was used as template to generate in vitro transcribed RNA which was then quantified and serially diluted (108 – 101 copies/μl).

## Acknowledgement

D.R.M. is funded by a Burroughs Wellcome Fund Postdoctoral Enrichment Program Award and a Hanna H. Gray Fellowship from the Howard Hughes Medical Institute and was supported by an <u>NIH</u> NIAID <u>T32 AI007151</u> and an NIH <u>F32 AI152296</u>. This work was supported by NIAID <u>R01 AI132178</u> and R01AI132178-04S1 (T.P.S. and R.S.B.) and an NIH animal models contract (<u>HHSN272201700036I</u>) to R.S.B. This project was supported in part by the North Carolina Policy Collaboratory at University of North Carolina at Chapel Hill with funding from the North Carolina Coronavirus Relief Fund established and appropriated by the North Carolina General Assembly.

Animal <u>histopathology</u> services were performed by the Animal Histopathology & Laboratory Medicine Core at the University of North Carolina, which is supported in part by an NCI Center Core Support Grant (<u>5P30CA016086-41</u>) to the UNC Lineberger Comprehensive Cancer Center.

