## Supplementary figures and images for "Therapeutic efficacy of an oral nucleoside analog of remdesivir against SARS-CoV-2 pathogenesis in mice"

### Supplemental Figure 1

Supplemental Figure 1

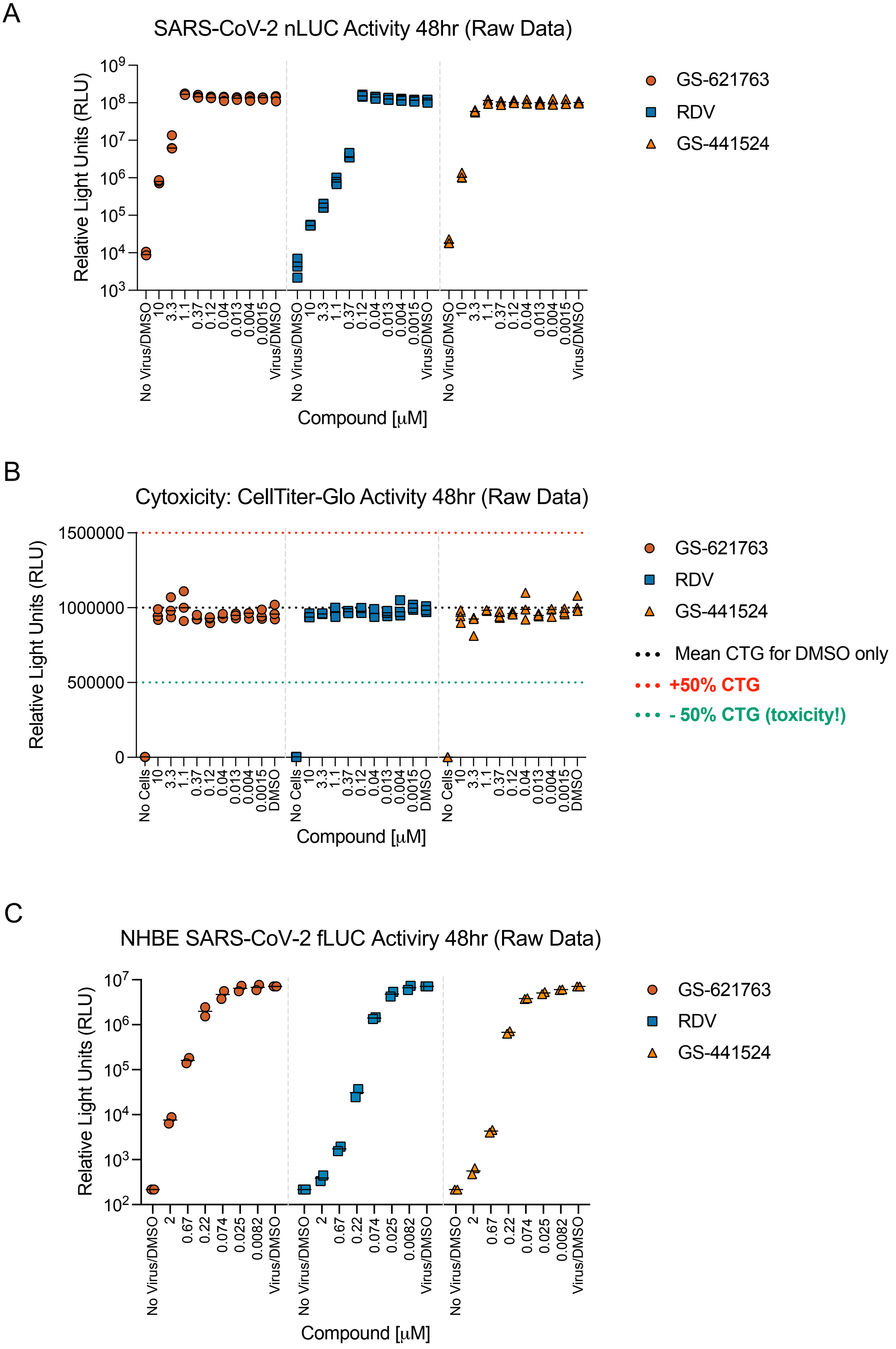

### Supplemental Figure 2

Supplemental Figure 2

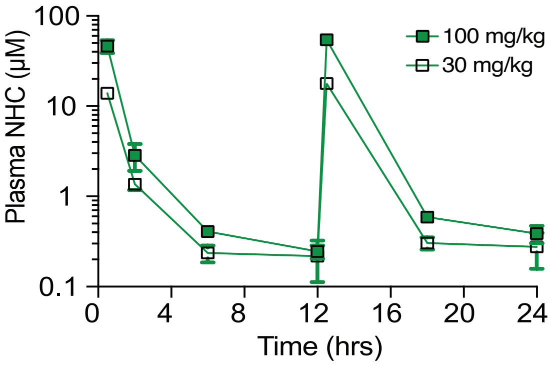
